# Hallmarks of tumor-experienced T cells are absent in multiple myeloma patients from diagnosis through maintenance therapy

**DOI:** 10.1101/2024.06.03.597178

**Authors:** Carolyn Shasha, David R. Glass, Ernest Moelhman, Laura Islas, Yuan Tian, Gregory L. Szeto, Tao Peng, Xiaoling Song, Michelle Wurscher, Thomas F. Bumol, Troy R. Torgerson, Philip D. Greenberg, Damian J. Green, Evan W. Newell

**Author notes:** Equal contribution (Carolyn Shasha, David R. Glass).

## Abstract

Dysregulation of the bone marrow (BM) niche in multiple myeloma (MM) alters the composition and state of resident immune cells, potentially impeding anti-tumor immunity. One common mechanism of immune inhibition in solid tumors is the induction of exhaustion in tumor-specific T cells. However, the extent of T cell tumor recognition and exhaustion is not well-characterized in MM. As the specific mechanisms of immune evasion are critical for devising effective therapeutic strategies, we deeply profiled the CD8^+^ T cell compartment of newly-diagnosed MM (NDMM) patients for evidence of tumor reactivity and T cell exhaustion. We applied single-cell multi-omic sequencing and antigen-specific mass cytometry to longitudinal BM and peripheral blood (PB) samples taken from timepoints spanning from diagnosis through induction therapy, autologous stem cell transplant (ASCT), and maintenance therapy. We identified an exhausted-like population that lacked several canonical exhaustion markers, was not significantly enriched in NDMM patients, and consisted of small, nonpersistent clones. We also observed an activated population with increased frequency in the PB of NDMM patients exhibiting phenotypic and clonal features consistent with homeostatic, antigen-nonspecific activation. However, there was no evidence of “tumor-experienced” T cells displaying hallmarks of terminal exhaustion and/or tumor-specific activation/expansion in NDMM patients at any timepoint.

## INTRODUCTION

MM is a hematological malignancy characterized by uncontrolled clonal expansion of plasma cells in the bone marrow. First-line treatment for NDMM is evolving, however induction therapy consisting of a three-drug combination including a proteasome inhibitor, immunomodulatory drug, and a steroid has been a standard of care.^1^ Eligible patients frequently proceed to ASCT (combined myeloablative melphalan chemotherapy and autologous stem cell rescue), followed by maintenance therapy (typically an immunomodulatory drug).

T cell exhaustion, characterized by diminished effector function and driven by chronic antigen stimulation such as from recognition of a persistent tumor,^2^ is a validated mechanism of immune evasion by many tumors.^3^ The immune microenvironment is already altered before diagnosis of active MM requiring therapy,^4^ but the presence of T cell exhaustion in NDMM and through first-line treatments has not previously been well-characterized. To explore tumor-specific recognition and exhaustion, we interrogated CD8^+^ T cells in NDMM patients, spanning diagnosis through induction, ASCT, and maintenance therapy.

## RESULTS

### Hallmarks of tumor-experienced T cells are absent in NDMM

We collected BM and PB samples from NDMM patients (n=9) before treatment (PRETX), during induction therapy (INDC), after induction therapy (EIND; median 33 days after start of last cycle), 90 days post-ASCT (TRN90), and 1-year post-ASCT (TRNY1; **Fig. 1A, Table S1**). We also collected HC samples (n=4 BM, n=4 PB) for comparison. To deeply profile the CD8^+^ T cell response and identify evidence of recent or ongoing tumor-antigen-specific T cell activity, we performed single-cell multi-omic sequencing and antigen-specific mass cytometry (**Table S2-5**). We clustered BM and PB T cells from MM patients and HCs into eleven distinct clusters based on their transcriptional and proteomic profiles (**Fig. 1B**). We identified two populations with atypical expression profiles: an activated cluster defined by the highest “activation” gene set score with genes such as *HLADRA* (HLA-DR) and *MKI67* (Ki-67, a proliferation marker), and an exhausted-like cluster defined by the highest “exhaustion” gene set score including inhibitory molecules such as *HAVCR2 (*TIM-3*), LAG3*, and *CTLA4* (**Fig. 1C-E**). Notably, neither cluster expressed canonical reactivity/exhaustion genes such as *PDCD1* (PD-1), *ENTPD1* (CD39), *TOX*, or *CXCL13* (**Fig. 1C**).^3^ As BM is predominantly the site of the tumor, we asked if any T cell populations were significantly enriched in BM as compared to PB. Only CD69^+^ tissue-resident memory (CD69 TRM) cells were enriched in the BM of MM patients, but this was also true in HCs (**Fig. S1A-B**). Activated and exhausted-like cells were not significantly differentially abundant between tissues.

**Figure 1:**
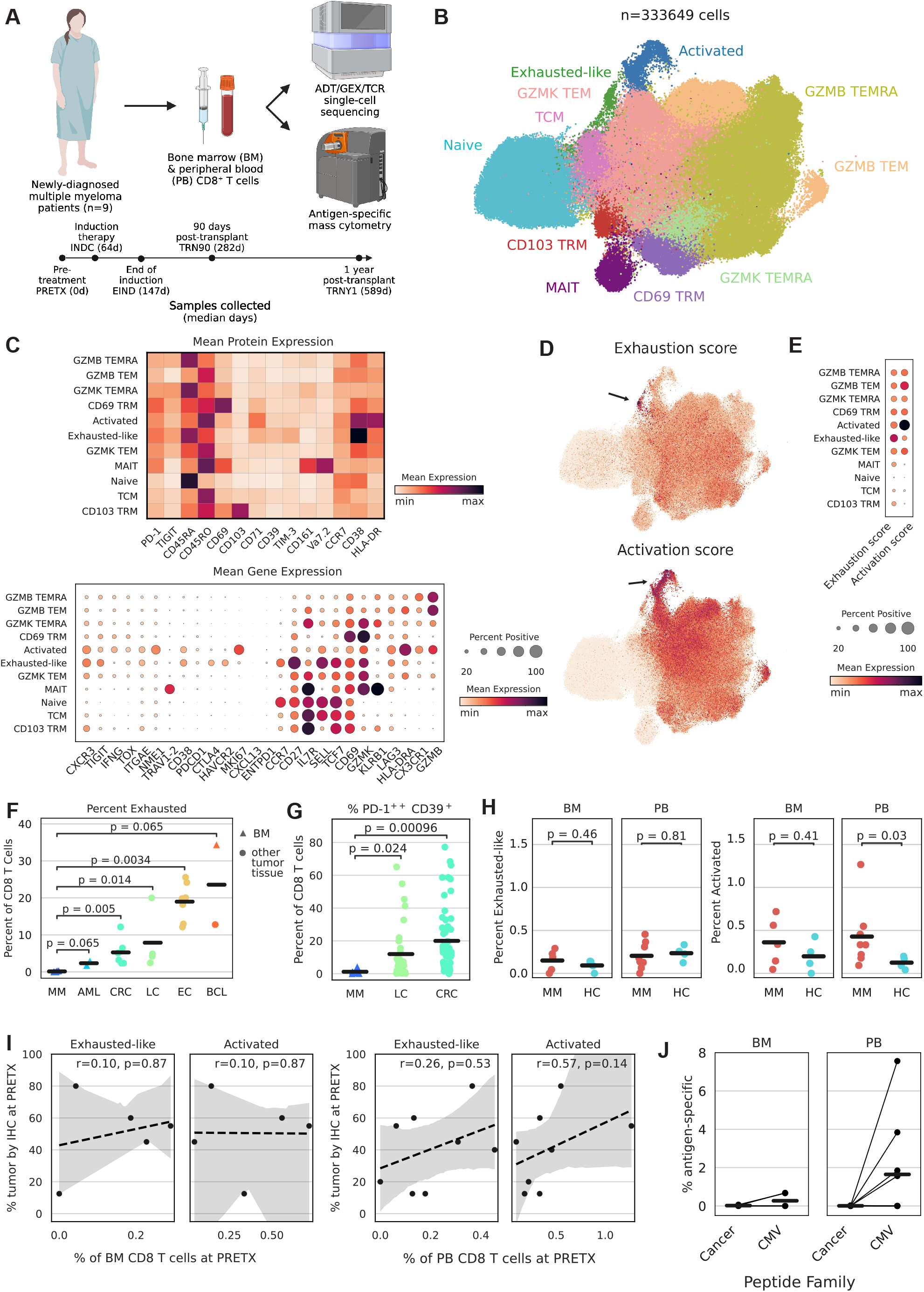
Hallmarks of tumor-experienced T cells are absent in NDMM. 1A: Experimental diagram and clinical time course. ADT=antibody-derived tag; GEX=gene expression; TCR=T cell receptor. 1B: TotalVI-UMAP of CD8^+^ T cells colored by cluster. 1C: Mean expression of proteins (top) or genes (bottom) by cluster. 1D: TotalVI-UMAP of CD8^+^ T cells colored by gene set score. 1E: Mean gene set score by cluster. 1F: PRETX frequency of exhausted-like/exhausted T cells quantified by single-cell sequencing, including public data,^5–8^ and separated by tumor type; p calculated by Wilcoxon rank sum test. AML=acute myeloid leukemia; CRC=colorectal cancer; LC=lung cancer; EC=esophageal cancer; BCL=B cell lymphoma. 1G: PRETX frequency of PD-1^++^CD39^+^ T cells quantified by mass cytometry, including public data,^9^ and separated by tumor type; p calculated by Wilcoxon rank sum test. 1H: Frequency of exhausted-like (left) or activated (right) cells in NDMM patients at PRETX and healthy controls, by tissue; p calculated by Wilcoxon rank sum test. 1I: Frequency of BM (left panel) or PB (right panel) exhausted-like or activated cells (x-axis) and BM tumor cells quantified by clinical immunohistochemistry (y-axis) at PRETX; r and p calculated by Spearman method. 1J: PRETX frequency of pMHC-tet^+^CD8^+^ T cells by peptide family and tissue; lines grouped by patient.

To compare the extent of exhaustion in MM to other cancer types, we integrated tumor-infiltrating CD8^+^ T cells from public datasets, including several from BM tumors, featured in a pan-cancer atlas.^5–8^ The frequency of exhausted-like cells in the BM of MM patients was lower than the frequency of exhausted cells in every other patient from every other cancer type (**Fig. 1F**). As an orthogonal confirmation, we manually gated exhausted-like cells as PD-1^++^CD39^+^ in our mass cytometry dataset and compared the frequency of these cells in the BM of MM patients to publicly available mass cytometry data from tumor-infiltrating CD8^+^ T cells in other cancer types.^9^ We found significantly fewer exhausted-like cells in MM samples compared to other tumors (**Fig. 1G, S1C**).

Activated and Granzyme B^+^ T effector memory (GZMB TEM) cells were significantly more abundant in the PB, but not BM, of PRETX samples from MM patients as compared to HCs, but no statistical differences were observed for exhausted-like cells (**Fig. 1H, S1D**). The frequency of PRETX BM tumor cells was not significantly correlated with the frequencies of activated or exhausted-like cells (**Fig. 1I**).

We also quantified antigen-specific T cells using an extensive panel of peptide-MHC tetramers (pMHC-tet) that included many tetramers presenting tumor antigens expressed in MM (**Table S3**), but we found no enrichment of tumor-specific T cells above background, despite routine detection of myeloma-unrelated, virus-specific T cells (**Fig. 1J, S2**). In summary, we observed no cell population expressing canonical markers of terminal exhaustion or tumor reactivity, and exhausted-like CD8^+^ T cells were significantly less abundant in NDMM compared to other cancer types and not significantly differentially abundant compared to HCs.

### Activated and exhausted-like cells lack clonal features typical of tumor-experienced T cells

Tumor cell frequency substantially diminished after induction therapy and ASCT (**Fig. 2A**). A tumor-reactive T cell population might follow these trends, but exhausted-like and activated T cells did not decrease in frequency over time and were significantly more abundant one-year post-ASCT than at PRETX in BM and PB, respectively (**Fig. 2B**). GZMB TEM cells were also significantly increased in frequency one-year post-ASCT in PB, while CD103 TRM cells were largely absent in BM and PB post-transplant (**Fig. S3A**). Tumor-reactive populations often exhibit increased levels of antigen-driven expansion of T cell receptor (TCR) clones,^3^ so we quantified the clonal composition of each cell population. The activated population contained a moderate proportion of expanded clones, significantly less than the GZMB TEM population but significantly more than the exhausted-like population, which had the smallest proportion and number of expanded clones across all populations (**Fig. 2C-D, S3B**). Antigen-specific clonal expansion often manifests as decreased TCR diversity in a cell population. At all timepoints, exhausted-like cells had higher mean TCR diversity than activated cells, which had higher mean TCR diversity than GZMB TEM cells (**Fig. S3C**). Activated and exhausted-like cells had few clones that significantly expanded or contracted over time, consistent with an absence of antigen-driven selection (**Fig. S3D**). To interrogate the specificities of clones in each cell population, we queried a database reporting experimentally-validated TCR-antigen pairs.^10^ We found activated and exhausted-like populations contained similar frequencies of bystander-specific clones to all other T cell populations, suggesting they are not uniquely enriched for tumor-specific clones (**Fig. 2E**).

**Figure 2:**
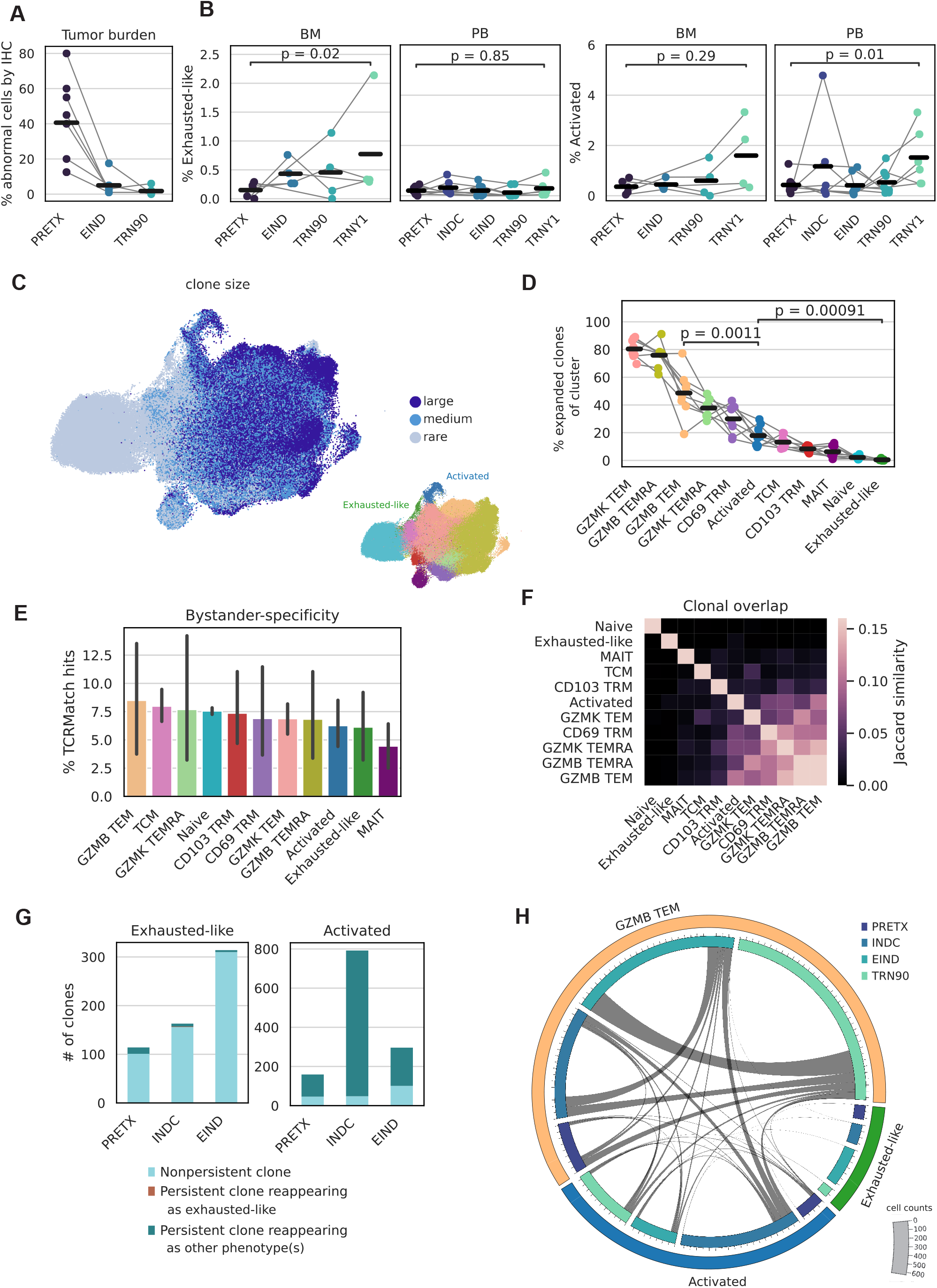
Activated and exhausted-like cells lack clonal features typical of tumor-experienced T cells. 2A: Frequency of BM tumor cells quantified by clinical immunohistochemistry, separated by timepoint; lines grouped by patient. 2B: Frequency of exhausted-like or activated cells by timepoint and tissue; lines grouped by patient; p calculated by Wilcoxon rank sum test. 2C: TotalVI-UMAP colored by clone size (left) or population (right). 2D: Frequency of expanded clones in NDMM patients by population; lines grouped by patient; p calculated by Wilcoxon rank sum test. 2E: Percent of bystander-specific clones in NDMM patients by population. 2F: Jaccard similarity of clonal composition in NDMM patients by population. 2G: Number of clones by population and timepoint, colored by clonal persistence at later timepoints. 2H: Circos plot by population and timepoint. Bar size indicates number of cells and lines indicate number of clonal connections.

Exhausted cells arise from non-exhausted precursors, which can manifest as an abundance of TCR clonotypes shared between phenotypes.^11^ However, exhausted-like cells had minimal clonal overlap with any other cell population (**Fig. 2F**). There was substantial clonal overlap between activated and GZMB TEM populations, even accounting for differences in cell counts (**Fig. S3E**), but the overlap was similar between MM patients and HCs (**Fig. S3F**), suggesting it was not driven by tumor antigen.

Chronic activation of tumor-specific T cells can result in accumulation of cells with an exhausted phenotype over time.^11^ However, activated clones never adopted an exhausted-like phenotype in subsequent timepoints, while exhausted-like clones infrequently appeared in subsequent timepoints; when they did, they rarely retained an exhausted-like phenotype (**Fig. 2G**). We observed many activated cells that were clonally related to GZMB TEM cells at subsequent timepoints and vice versa, consistent with antigen-nonspecific homeostatic proliferative bursting (**Fig. 2H**). In summary, exhausted-like cells were comprised of non-expanded clones that rarely appeared at multiple timepoints, while activated cells displayed substantial clonal overlap with GZMB TEM cells, consistent with homeostatic activation and a lack of antigen-driven selection.

## DISCUSSION

Other groups have reported a paucity of myeloma-responsive T cells^12^ and myeloma-specific TCRs^13^ in NDMM. Here, we report the absence of transcriptional, proteomic, and clonal hallmarks of tumor-experienced CD8^+^ T cells from NDMM patients, from diagnosis through maintenance therapy. Our findings suggest myeloma cell escape from T cell killing in NDMM is primarily mediated through evasion of recognition by antigen-specific T cells, rather than through induction of exhaustion in an established, clonally-expanded, tumor-specific T cell population. Activated cells were significantly enriched in the PB of MM patients compared to HCs but displayed evidence of homeostatic activation rather than tumor antigen-specific activation. Other groups have noted increased abundances of T-helper 17 cells,^14^ regulatory T cells,^15^ and TIGIT^+^CD8^+^ T cells^16^ in NDMM patients, concordant with the idea that MM induces global immunomodulatory effects on the T cell compartment that may limit antigen recognition. It is possible that T cell exhaustion plays a role in a minority of NDMM cases not revealed by the sampling here. One international staging system stage III NDMM patient from a pan-cancer atlas was reported to have detectable levels of exhausted T cells,^8,17^ so the frequency of this occurrence and the associated disease features could be investigated. Anti-SOX2 immunity was observed in patients with monoclonal gammopathy^18^ and in MM patients after allotransplant^19^ and CAR-T therapy,^20^ so the phenotypes of SOX2-specific T cells could be investigated for evidence of exhaustion in these contexts.

Checkpoint blockade monotherapy has performed poorly in refractory and/or relapsed MM (RRMM).^21^ In solid tumors, evidence of pre-existing, tumor-reactive T cells predicts response to checkpoint blockade,^22,23^ while patients with cold tumors lacking tumor-reactive T cells tend to respond poorly.^24^ While our study did not include samples from RRMM patients, our data suggest that combination immunotherapies that draw inspiration from strategies targeting cold solid tumors^25^ may enhance T cell participation in the anti-myeloma response and increase therapeutic efficacy.

## METHODS

### Clinical cohort and samples

NDMM patients diagnosed and treated at the Fred Hutchinson Cancer Center and HCs provided written informed consent, and analysis was performed as approved by FHIRB10265. NDMM patients received induction therapy, stem cell mobilization, melphalan, ASCT, and maintenance therapy (**Fig. 1A, Table S1**). Longitudinal BM and PB samples were collected from diagnosis through maintenance therapy. Mononuclear cells were isolated by density gradient separation, cryopreserved, and stored in liquid nitrogen.

### Mass cytometry antibody conjugation and pMHC tetramer generation

Purified antibodies lacking carrier proteins were purchased (**Table S2**), and antibody conjugation was performed according to the manufacturer”s protocol (Standard Biotools), as previously described.^26^ Peptides for detection of antigen-specific responses were selected to encompass a wide range of myeloma, cancer, and bystander targets (**Table S3**). Common HLA alleles were chosen for antigen-specific reagents, as patients were not HLA-typed. Tetramer generation and conjugation was performed as previously described.^27^ Briefly, a unique code was generated by conjugating streptavidin to three metal isotopes as above and each pMHC was associated with a specific three metal combination.^28^ For each pMHC, 5 mL peptide (1 mM) was added to 100 mL HLA monomer (100 mg/mL, diluted in PBS) and exposed to UV (365 nm) for 10 min for peptide exchange and left overnight at 4C. For tetramerization, 5 mL of labeled streptavidin (50 mg/mL) was mixed with pMHC and incubated for 10 min at 25C. This was repeated three additional times for a total addition of 20 mL. Then, tetramers were incubated with free biotin (10 mM) for 10 min. Tetramers were combined and concentrated using a 50-kDa Amicon filter (Milipore). Antibody and tetramer cocktails were cryopreserved in cell-staining media (CSM; PBS, 0.5% BSA, 0.02% sodium azide) at -80C.

### Mass cytometry staining and acquisition

Samples were processed in batches with common healthy PBMC samples control samples included for assessment of batch effect. Cells were washed in CSM and stained for cisplatin (5 mM) in PBS for 5 min at 25C. Cells were washed and stained with tetramer cocktails in CSM for 1 hr at 25C. Cells were washed, stained with primary antibody cocktails for 25 min at 4C, washed, stained with secondary surface antibody cocktails for 25 min at 4C, and fixed overnight in 2% paraformaldehyde. Cells were washed, permeabilized using permeabilization buffer according to manufacturer”s instructions (eBioscience), washed, and stained with intracellular antibodies for 30 min at 25C. Cells were washed and incubated with DNA intercalator (Standard Biotools) for 10 min at 25C. Cells were washed three times with Cell Acquisition Solution (Standard Biotools) and run on the Helios mass cytometer (Standard Biotools).

### Mass cytometry data processing and analysis

Fcs files were bead normalized^29^ and compensated.^30^ Zero values were randomized with a uniform distribution between zero and -1. Files were hand-gated in FlowJo to viable CD8^+^ T cell singlets (**Fig. S4A**) and uploaded to cellengine.com. pMHC tetramers and antibodies were hand-gated to account for batch effect. Cancer pMHC tetramers were background subtracted using the healthy normalization control samples for each batch. Public data^9^ were already gated to viable CD8^+^ T cell singlets by the original authors, so those fcs files were uploaded to cellengine.com and hand-gated alongside the original data generated in this study.

### Single-cell sequencing data acquisition

pMHC monomers (BioLegend) were generated and peptide-exchanged as above. Monomers were tetramerized with TotalSeq-C PE streptavidin following manufacturer”s instructions. Samples were thawed and stained with Human TruStain FcX (BioLegend) for 10 min at 25C, washed, then stained with tetramer cocktail for 1 hour at 25C. After washing, cells were stained with fluorescent antibodies (**Table S4**) and TotalSeq-C antibodies (**Table S5**), which had been combined and filtered using a 0.1mm filter (Millipore). Cells were washed two more times in CSM, then CD8^+^ T cells were isolated by fluorescence-activated cell sorting (**Fig. S4B**). Some enrichment for pMHC-tet^+^ cells and CD69^+^ cells (BM only) was performed but this had negligible effects on total sorted cell counts. Sorted cells were subjected to 10x library preparation following manufacturer”s instructions (User Guide: CG000424) and loaded onto the 10x Chromium controller using Chromium Next GEM Single Cell 5” Reagent Kits v2 (Dual Index) with Feature Barcoding technology for Cell-Surface Protein and Immune Receptor Mapping (10x Genomics).

### Single-cell sequencing data preprocessing

Raw RNA data and antibody-derived tag sequences were demultiplexed and aligned to the GRCh38 reference genome using Cell Ranger v6.0.1 (10x Genomics). T cell receptor (TCR) sequences were also identified using Cell Ranger v6.0.1. The filtered expression matrices (Cell Ranger outputs) were pre-processed using Scanpy^31^ v1.7.2 in Python. Low quality cells with fewer than 200 or more than 4000 expressed genes were removed, and cells with fewer than 200 counts or more than 25% mitochondrial gene expression were removed. Doublets were removed with Scrublet^32^ v0.2.1 using its automatic doublet detection threshold. TCR sequences were read in and processed using Scirpy^33^ v0.10.1. Cells lacking an immune receptor as measured by TCR sequencing were removed as well. CD8^+^ T cells were extracted by identifying CD3^+^, CD8^+^, and CD4^-^ cells based on ADT expression.

Integration of samples (patients, healthy donors, and tissue type/timepoint) was performed using totalVI^34^ v0.17.0, which integrates datasets using both gene and ADT expression. To run totalVI, the top 3000 highly variable genes were used (identified using the “seurat_v3” method in Scanpy). Mitochondrial and ribosomal genes were then removed. The variational autoencoder model was trained with 250 max epochs, with all other parameters set to defaults. The resulting latent representation was used to generate a nearest-neighbors map and UMAP representation. ADT expression was transformed using CLR normalization, and gene expression was log-normalized.

### Single-cell sequencing data analysis

Unsupervised Leiden clustering was performed in Scanpy using the totalVI-generated neighbors graph. Clusters were labeled according to canonical gene and protein markers (**Fig. 1C**). Within broad cell types (TEM, TEMRA, and TRM) where multiple clusters were identified, clusters were labeled descriptively based on what we identified as the most prominent gene/protein expression difference among the clusters (i.e. GZMB^+^ vs. GZMK^+^ TEM/TEMRA, and CD69^+^ vs. CD103^+^ TRM).

Common exhaustion genes (*CTLA4, HAVCR2, LAG3, PDCD1, TIGIT*, and *ENTPD1*) and activation/proliferation genes (*NME1, IL2RA, CD38, HLA-DRA, HLA-DRB1, HLA-DRB3, HLA-DRB4, HLA-DRB5, MKI67*, and *PCNA*) were used to calculate the exhaustion and activation gene scores, respectively. Gene scores were calculated using the Scanpy “score_genes” function.

External data for solid tumor comparisons were downloaded from the link (http://cancer-pku.cn:3838/PanC_T/) provided by Zheng et al.^8^ Raw expression data for CD8^+^ T cells were downloaded per dataset. Following the same steps as our in-house data, the data was log-normalized and gene scores were calculated using the Scanpy “score_genes” function.

Clonal analysis (e.g. calculation of Jaccard similarity and clonal diversity) was performed using the Scirpy package. Scipy^35^ v1.10.1 (Virtanen et al 2020) was used to perform statistical correlations, and the statannot v0.2.3 package was used to add statistical annotations to plots.

## ACKNOWLEDGEMENTS

We thank Simone Minnie, Koshlan Mayer-Blackwell, and Lucas Graybuck for helpful discussions. We also thank Christine Crider and the clinical and administrative teams for their support as well as Mark Majeres for data management. D.R.G. was supported by a Cancer Research Institute Irvington Postdoctoral Fellowship and an American Society of Hematology Fellow Scholar Award. X.S. is partially supported by the Institute of Translational Health Sciences, which is funded by the National Center for Advancing Translational Sciences of the National Institutes of Health under award number UL1TR002319. P.D.G. was supported in part by grants from the Allen Institute of Immunology, the Parker Institute for Cancer Immunotherapy, and the NCI: CA18029. This work was supported by Fred Hutchinson Cancer Center New Development funds (E.W.N.), The Any Hill Endowment Distinguished Researcher CARE fund (E.W.N), Defeat Myeloma, and the Allen Institute for Immunology.

## AUTHORSHIP CONTRIBUTIONS

Conceptualization, C.S., D.R.G., T.F.B, T.R.T., P.D.G., D.J.G., E.W.N.; Methodology, C.S., D.R.G., Y.T., G.L.S., T.P., E.W.N.; Investigation, E.M., L.I.; Formal analysis, C.S., D.R.G.; Data curation, C.S., D.R.G., X.S., M.W.; Writing – Original Draft, C.S., D.R.G., E.M.; Writing – Review & Editing, C.S., D.R.G., G.L.S., P.D.G., D.J.G., E.W.N.; Funding Acquisition, T.F.B., T.R.T., P.D.G., D.J.G., E.W.N.; Resources, T.F.B., T.R.T., P.D.G., D.J.G., E.W.N.; Supervision, T.F.B., T.R.T., P.D.G., D.J.G., E.W.N.

## REFERENCES

1. Cowan AJ, Green DJ, Kwok M, et al. Diagnosis and Management of Multiple Myeloma: A Review. JAMA. 2022;327(5):464–477.

2. Dolina JS, Van Braeckel-Budimir N, Thomas GD, Salek-Ardakani S. CD8+ T Cell Exhaustion in Cancer. Front. Immunol. 2021;12:.

3. Chow A, Perica K, Klebanoff CA, Wolchok JD. Clinical implications of T cell exhaustion for cancer immunotherapy. Nat Rev Clin Oncol. 2022;19(12):775–790.

4. García-Ortiz A, Rodríguez-García Y, Encinas J, et al. The Role of Tumor Microenvironment in Multiple Myeloma Development and Progression. Cancers. 2021;13(2):217.

5. van Galen P, Hovestadt V, Wadsworth II MH, et al. Single-Cell RNA-Seq Reveals AML Hierarchies Relevant to Disease Progression and Immunity. Cell. 2019;176(6):1265-1281.e24.

6. Song Q, Hawkins GA, Wudel L, et al. Dissecting intratumoral myeloid cell plasticity by single cell RNA-seq. Cancer Medicine. 2019;8(6):3072–3085.

7. Zhang L, Yu X, Zheng L, et al. Lineage tracking reveals dynamic relationships of T cells in colorectal cancer. Nature. 2018;564(7735):268–272.

8. Zheng L, Qin S, Si W, et al. Pan-cancer single-cell landscape of tumor-infiltrating T cells. Science. 2021;374(6574):abe6474.

9. Simoni Y, Becht E, Fehlings M, et al. Bystander CD8 + T cells are abundant and phenotypically distinct in human tumour infiltrates. Nature. 2018;557(7706):575–579.

10. Goncharov M, Bagaev D, Shcherbinin D, et al. VDJdb in the pandemic era: a compendium of T cell receptors specific for SARS-CoV-2. Nat Methods. 2022;19(9):1017–1019.

11. Philip M, Schietinger A. CD8+ T cell differentiation and dysfunction in cancer. Nat Rev Immunol. 2022;22(4):209–223.

12. Dhodapkar MV, Krasovsky J, Olson K. T cells from the tumor microenvironment of patients with progressive myeloma can generate strong, tumor-specific cytolytic responses to autologous, tumor-loaded dendritic cells. Proceedings of the National Academy of Sciences. 2002;99(20):13009–13013.

13. Welters C, Lammoglia Cobo MF, Stein CA, et al. Immune Phenotypes and Target Antigens of Clonally Expanded Bone Marrow T Cells in Treatment-Naïve Multiple Myeloma. Cancer Immunology Research. 2022;10(11):1407–1419.

14. Prabhala RH, Pelluru D, Fulciniti M, et al. Elevated IL-17 produced by Th17 cells promotes myeloma cell growth and inhibits immune function in multiple myeloma. Blood. 2010;115(26):5385–5392.

15. Giannopoulos K, Kaminska W, Hus I, Dmoszynska A. The frequency of T regulatory cells modulates the survival of multiple myeloma patients: detailed characterisation of immune status in multiple myeloma. Br J Cancer. 2012;106(3):546–552.

16. Guillerey C, Harjunpää H, Carrié N, et al. TIGIT immune checkpoint blockade restores CD8+ T-cell immunity against multiple myeloma. Blood. 2018;132(16):1689–1694.

17. Minnie SA, Waltner OG, Zhang P, et al. TIM-3+ CD8 T cells with a terminally exhausted phenotype retain functional capacity in hematological malignancies. Science Immunology. 2024;9(94):eadg1094.

18. Spisek R, Kukreja A, Chen L-C, et al. Frequent and specific immunity to the embryonal stem cell–associated antigen SOX2 in patients with monoclonal gammopathy. Journal of Experimental Medicine. 2007;204(4):831–840.

19. Kobold S, Tams S, Luetkens T, et al. Patients with multiple myeloma develop SOX2-specific autoantibodies after allogeneic stem cell transplantation. Clin Dev Immunol. 2011;2011:302145.

20. Garfall AL, Stadtmauer EA, Hwang W-T, et al. Anti-CD19 CAR T cells with high-dose melphalan and autologous stem cell transplantation for refractory multiple myeloma. JCI Insight. 2019;3(8):.

21. Abramson HN. Immunotherapy of Multiple Myeloma: Current Status as Prologue to the Future. International Journal of Molecular Sciences. 2023;24(21):15674.

22. Thommen DS, Koelzer VH, Herzig P, et al. A transcriptionally and functionally distinct PD-1+ CD8+ T cell pool with predictive potential in non-small-cell lung cancer treated with PD-1 blockade. Nat Med. 2018;24(7):994–1004.

23. Au L, Hatipoglu E, Massy MR de, et al. Determinants of anti-PD-1 response and resistance in clear cell renal cell carcinoma. Cancer Cell. 2021;39(11):1497-1518.e11.

24. Herbst RS, Soria J-C, Kowanetz M, et al. Predictive correlates of response to the anti-PD-L1 antibody MPDL3280A in cancer patients. Nature. 2014;515(7528):563–567.

25. Bonaventura P, Shekarian T, Alcazer V, et al. Cold Tumors: A Therapeutic Challenge for Immunotherapy. Front. Immunol. 2019;10:.

26. Hartmann FJ, Simonds EF, Vivanco N, et al. Scalable Conjugation and Characterization of Immunoglobulins with Stable Mass Isotope Reporters for Single-Cell Mass Cytometry Analysis. Mass Cytometry: Methods and Protocols. 2019;55–81.

27. Simoni Y, Fehlings M, Newell EW. Multiplex MHC Class I Tetramer Combined with Intranuclear Staining by Mass Cytometry. Mass Cytometry: Methods and Protocols. 2019;147–158.

28. Newell EW, Sigal N, Nair N, et al. Combinatorial tetramer staining and mass cytometry analysis facilitate T-cell epitope mapping and characterization. Nat Biotechnol. 2013;31(7):623–629.

29. Finck R, Simonds EF, Jager A, et al. Normalization of mass cytometry data with bead standards. Cytometry Part A. 2013;83A(5):483–494.

30. Chevrier S, Crowell HL, Zanotelli VRT, et al. Compensation of Signal Spillover in Suspension and Imaging Mass Cytometry. Cell Systems. 2018;6(5):612-620.e5.

31. Wolf FA, Angerer P, Theis FJ. SCANPY: large-scale single-cell gene expression data analysis. Genome Biology. 2018;19(1):15.

32. Wolock SL, Lopez R, Klein AM. Scrublet: Computational Identification of Cell Doublets in Single-Cell Transcriptomic Data. cels. 2019;8(4):281-291.e9.

33. Sturm G, Szabo T, Fotakis G, et al. Scirpy: a Scanpy extension for analyzing single-cell T-cell receptor-sequencing data. Bioinformatics. 2020;36(18):4817–4818.

34. Gayoso A, Steier Z, Lopez R, et al. Joint probabilistic modeling of single-cell multi-omic data with totalVI. Nat Methods. 2021;18(3):272–282.

35. Virtanen P, Gommers R, Oliphant TE, et al. SciPy 1.0: fundamental algorithms for scientific computing in Python. Nat Methods. 2020;17(3):261–272.

