## Supplemental figures for "Hallmarks of tumor-experienced T cells are absent in multiple myeloma patients from diagnosis through maintenance therapy"

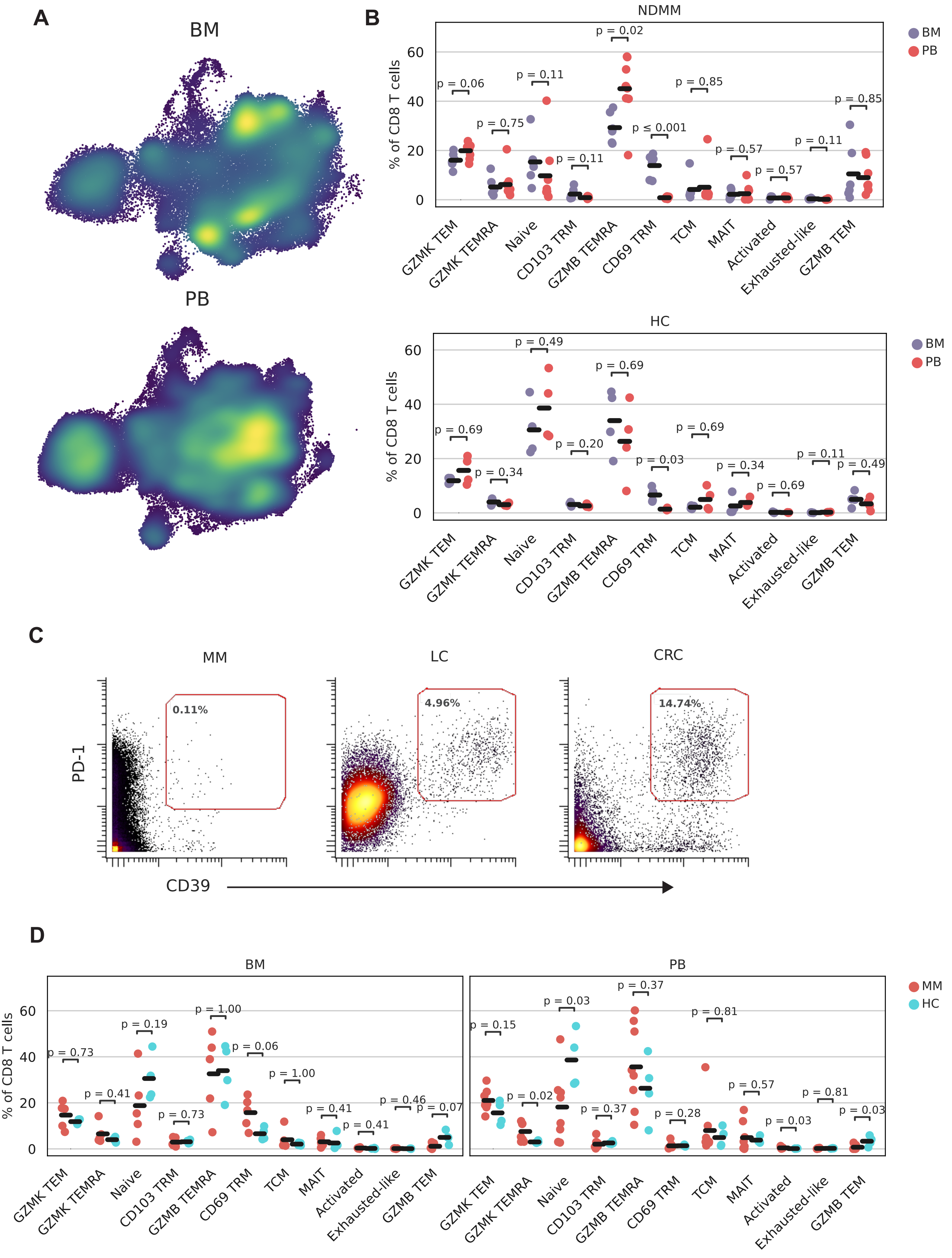

A
BM
PB
B
NDMM
HC
BM
PB
% of CD8 T cells
GZMK TEM
GZMK TEMRA
Naive
CD103 TRM
GZMB TEMRA
CD69 TRM
TCM
MAIT
Activated
Exhausted-like
GZMB TEM
C
MM
LC
CRC
0.11%
4.96%
14.74%
PD-1
CD39
D
MM
HC

**Figure S1: Cell population and antigen-specificity quantification**
S1A: TotalVI-UMAP separated by tissue and colored by density
S1B: Frequency of cell populations by tissue, separated into newly-diagnosed multiple myeloma (NDMM) patients (top) and HCs (bottom); p calculated by Wilcoxon rank sum test.
S1C: Representative gating of PD-$1^{++}$CD$39^{+}$ CD$8^{+}$ T cells from NDMM, lung cancer (LC), and colorectal cancer (CRC) patients.
S1D: Frequency of cell populations by disease status (MM or HC), separated by tissue; p calculated by Wilcoxon rank sum test.

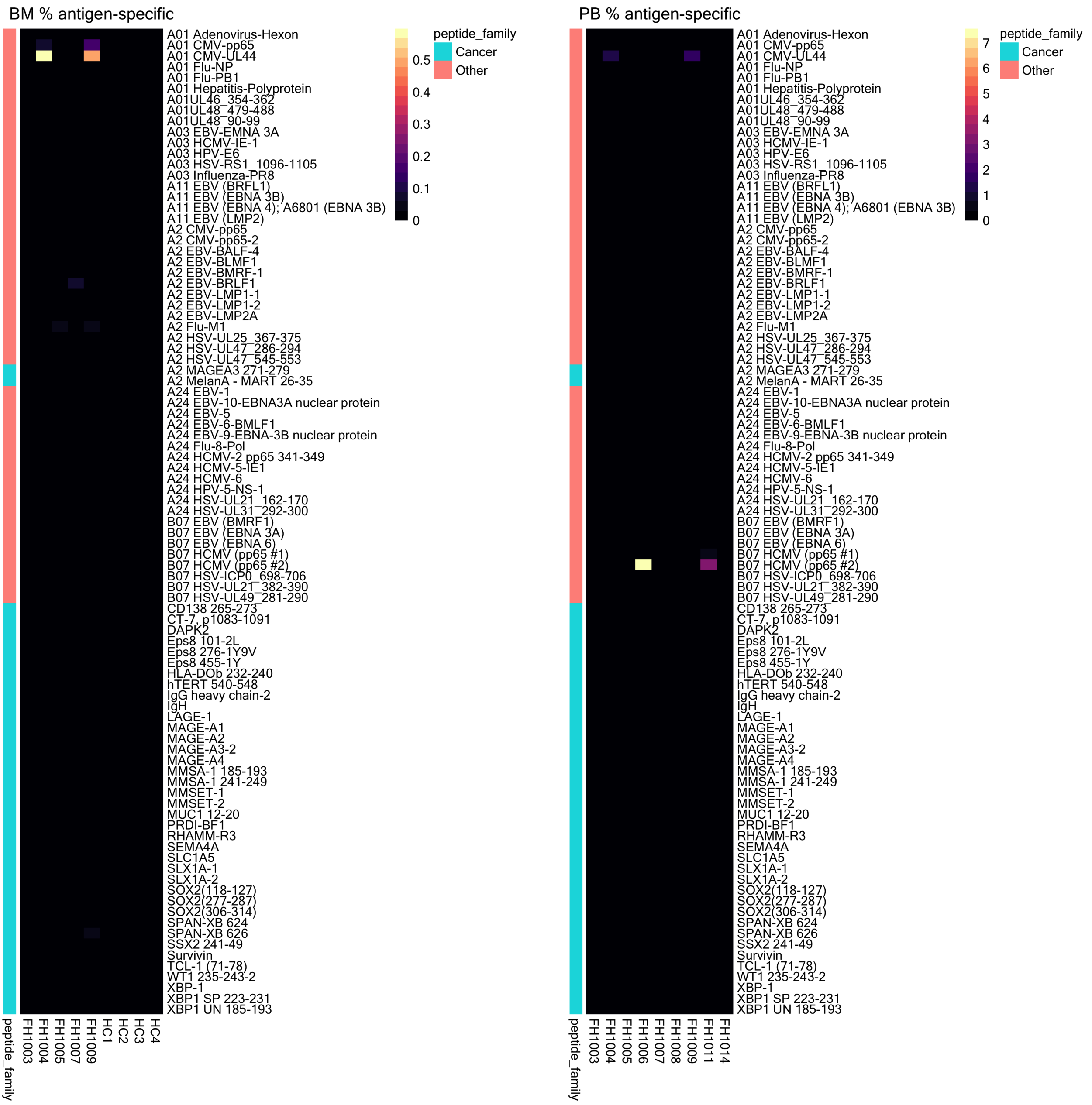
BM % antigen-specific
A01 Adenovirus-Hexon
A01 CMV-pp65
A01 CMV-UL44
A01 Flu-NP
A01 Flu-PB1
A01 Hepatitis-Polyprotein
A01UL46_354-362
A01UL48_479-488
A01UL48_90-99
A03 EBV-EMNA 3A
A03 HCMV-IE-1
A03 HPV-E6
A03 HSV-RS1_1096-1105
A03 Influenza-PR8
A11 EBV (BRFL1)
A11 EBV (EBNA 3B)
A11 EBV (EBNA 4); A6801 (EBNA 3B)
A11 EBV (LMP2)
A2 CMV-pp65
A2 CMV-pp65-2
A2 EBV-BALF-4
A2 EBV-BLMF1
A2 EBV-BMRF-1
A2 EBV-BRLF1
A2 EBV-LMP1-1
A2 EBV-LMP1-2
A2 EBV-LMP2A
A2 Flu-M1
A2 HSV-UL25_367-375
A2 HSV-UL47_286-294
A2 HSV-UL47_545-553
A2 MAGEA3 271-279
A2 MelanA - MART 26-35
A24 EBV-1
A24 EBV-10-EBNA3A nuclear protein
A24 EBV-5
A24 EBV-6-BMLF1
A24 EBV-9-EBNA-3B nuclear protein
A24 Flu-8-Pol
A24 HCMV-2 pp65 341-349
A24 HCMV-5-IE1
A24 HCMV-6
A24 HPV-5-NS-1
A24 HSV-UL21_162-170
A24 HSV-UL31_292-300
B07 EBV (BMRF1)
B07 EBV (EBNA 3A)
B07 EBV (EBNA 6)
B07 HCMV (pp65 #1)
B07 HCMV (pp65 #2)
B07 HSV-ICP0_698-706
B07 HSV-UL21_382-390
B07 HSV-UL49_281-290
CD138 265-273
CT-7, p1083-1091
DAPK2
Eps8 101-2L
Eps8 276-1Y9V
Eps8 455-1Y
HLA-DOb 232-240
hTERT 540-548
IgG heavy chain-2
IgH
LAGE-1
MAGE-A1
MAGE-A2
MAGE-A3-2
MAGE-A4
MMSA-1 185-193
MMSA-1 241-249
MMSET-1
MMSET-2
MUC1 12-20
PRDI-BF1
RHAMM-R3
SEMA4A
SLC1A5
SLX1A-1
SLX1A-2
SOX2(118-127)
SOX2(277-287)
SOX2(306-314)
SPAN-XB 624
SPAN-XB 626
SSX2 241-49
Survivin
TCL-1 (71-78)
WT1 235-243-2
XBP-1
XBP1 SP 223-231
XBP1 UN 185-193
peptide_family
FH1003
FH1004
FH1005
FH1007
FH1009
HC1
HC2
HC3
HC4
0.5
0.4
0.3
0.2
0.1
0
peptide_family
Cancer
Other
PB % antigen-specific
A01 Adenovirus-Hexon
A01 CMV-pp65
A01 CMV-UL44
A01 Flu-NP
A01 Flu-PB1
A01 Hepatitis-Polyprotein
A01UL46_354-362
A01UL48_479-488
A01UL48_90-99
A03 EBV-EMNA 3A
A03 HCMV-IE-1
A03 HPV-E6
A03 HSV-RS1_1096-1105
A03 Influenza-PR8
A11 EBV (BRFL1)
A11 EBV (EBNA 3B)
A11 EBV (EBNA 4); A6801 (EBNA 3B)
A11 EBV (LMP2)
A2 CMV-pp65
A2 CMV-pp65-2
A2 EBV-BALF-4
A2 EBV-BLMF1
A2 EBV-BMRF-1
A2 EBV-BRLF1
A2 EBV-LMP1-1
A2 EBV-LMP1-2
A2 EBV-LMP2A
A2 Flu-M1
A2 HSV-UL25_367-375
A2 HSV-UL47_286-294
A2 HSV-UL47_545-553
A2 MAGEA3 271-279
A2 MelanA - MART 26-35
A24 EBV-1
A24 EBV-10-EBNA3A nuclear protein
A24 EBV-5
A24 EBV-6-BMLF1
A24 EBV-9-EBNA-3B nuclear protein
A24 Flu-8-Pol
A24 HCMV-2 pp65 341-349
A24 HCMV-5-IE1
A24 HCMV-6
A24 HPV-5-NS-1
A24 HSV-UL21_162-170
A24 HSV-UL31_292-300
B07 EBV (BMRF1)
B07 EBV (EBNA 3A)
B07 EBV (EBNA 6)
B07 HCMV (pp65 #1)
B07 HCMV (pp65 #2)
B07 HSV-ICP0_698-706
B07 HSV-UL21_382-390
B07 HSV-UL49_281-290
CD138 265-273
CT-7, p1083-1091
DAPK2
Eps8 101-2L
Eps8 276-1Y9V
Eps8 455-1Y
HLA-DOb 232-240
hTERT 540-548
IgG heavy chain-2
IgH
LAGE-1
MAGE-A1
MAGE-A2
MAGE-A3-2
MAGE-A4
MMSA-1 185-193
MMSA-1 241-249
MMSET-1
MMSET-2
MUC1 12-20
PRDI-BF1
RHAMM-R3
SEMA4A
SLC1A5
SLX1A-1
SLX1A-2
SOX2(118-127)
SOX2(277-287)
SOX2(306-314)
SPAN-XB 624
SPAN-XB 626
SSX2 241-49
Survivin
TCL-1 (71-78)
WT1 235-243-2
XBP-1
XBP1 SP 223-231
XBP1 UN 185-193
peptide_family
FH1003
FH1004
FH1005
FH1006
FH1007
FH1008
FH1009
FH1011
FH1014
7
6
5
4
3
2
1
0
peptide_family
Cancer
Other

**Figure S2: PRETX Frequency of peptide-MHC tetramer$^+$CD8$^+$ T cells by peptide, tissue, and individual.**

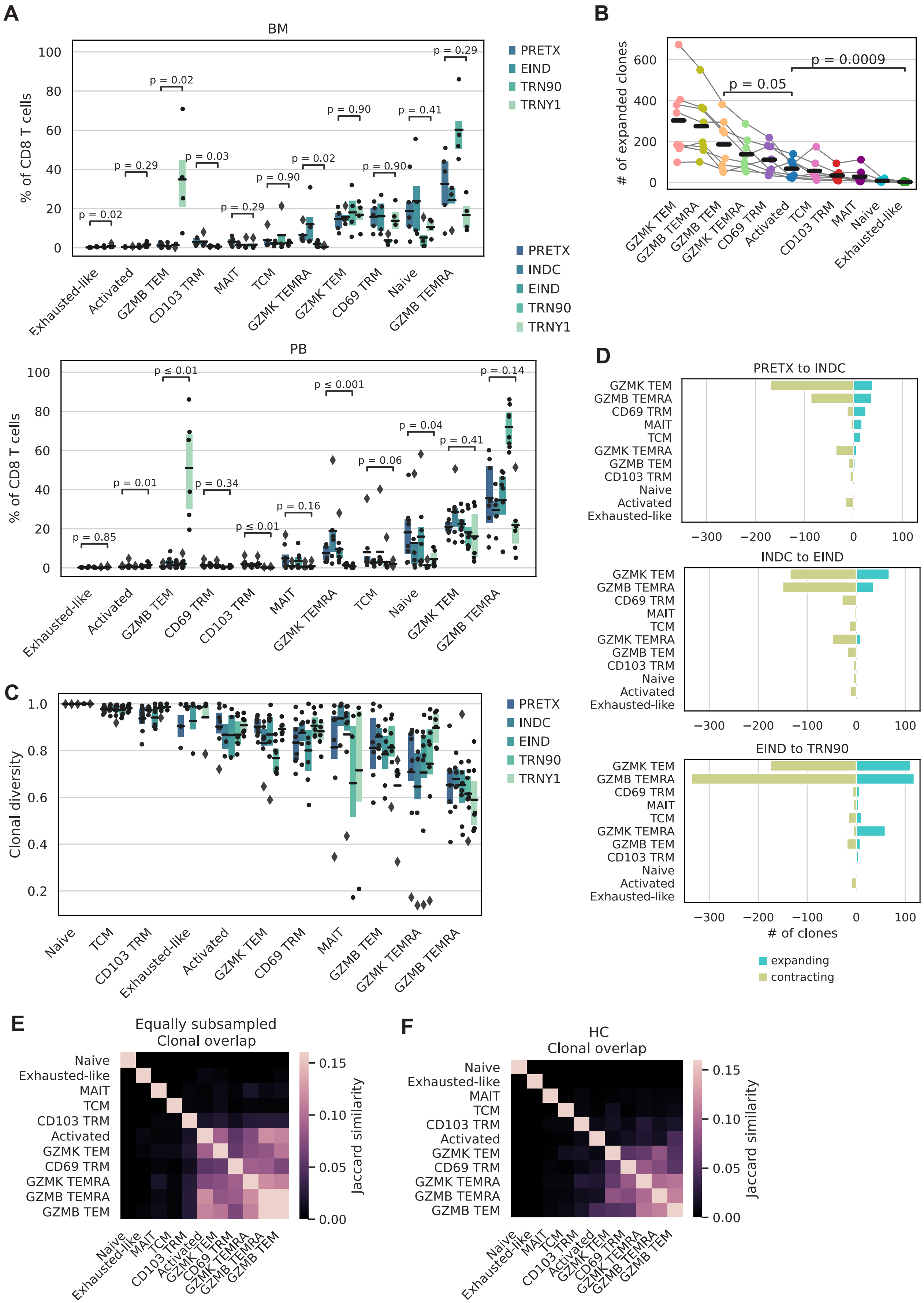

A
BM
% of CD8 T cells
PRETX
EIND
TRN90
TRNY1
PRETX
INDC
EIND
TRN90
TRNY1
Exhausted-like
Activated
GZMB TEM
CD103 TRM
MAIT
TCM
GZMK TEMRA
GZMK TEM
CD69 TRM
Naive
GZMB TEMRA
PB
p ≤ 0.01
p ≤ 0.001
B
### of expanded clones
p = 0.05
p = 0.0009
C
Clonal diversity
D
PRETX to INDC
INDC to EIND
EIND to TRN90
### of clones
expanding
contracting
E
Equally subsampled
Clonal overlap
Jaccard similarity
F
HC
Clonal overlap
Jaccard similarity

**Figure S3: Clonal and temporal dynamics**

S3A: Frequency of cell populations by time and tissue; p calculated by Wilcoxon rank sum test.
S3B: Number of expanded clones by population; p calculated by Wilcoxon rank sum test.
S3C: Alpha diversity (Shannon entropy) of clones by time and population.
S3D: Number of significantly ($p<0.05$) expanding and contracting clones by population and timepoint; p calculated by Fisher's exact test.
S3E: Jaccard similarity of clonal composition by population, equally sampled by population.
S3F: Jaccard similarity of clonal composition by population, for HCs only

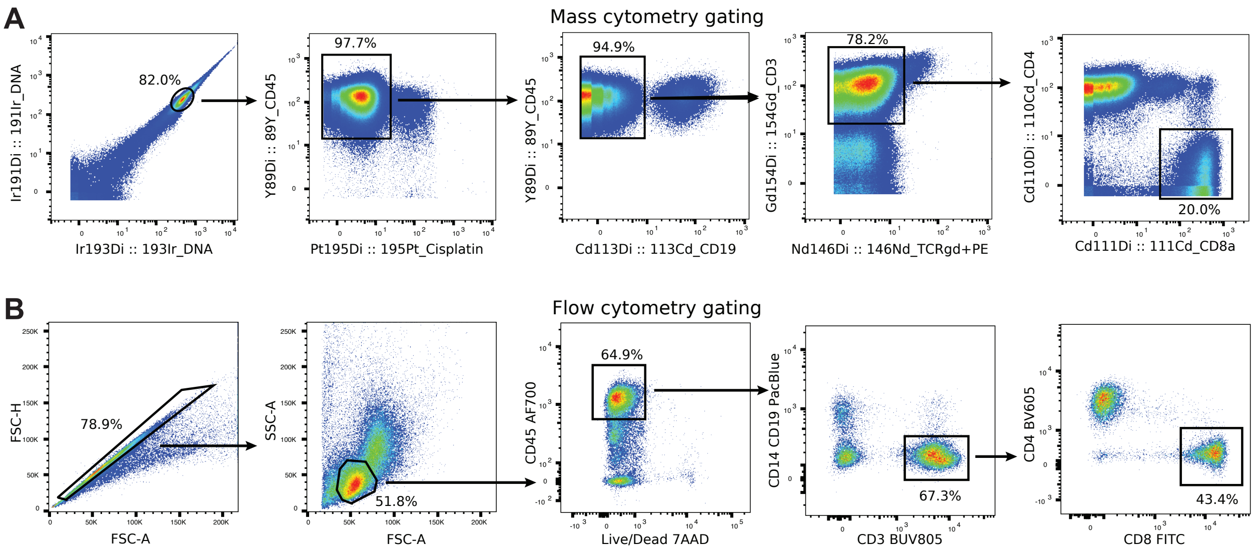

A
Mass cytometry gating
82.0%
Ir191Di :: 191Ir_DNA
Ir193Di :: 193Ir_DNA
97.7%
Y89Di :: 89Y_CD45
Pt195Di :: 195Pt_Cisplatin
94.9%
Y89Di :: 89Y_CD45
Cd113Di :: 113Cd_CD19
78.2%
Gd154Di :: 154Gd_CD3
Nd146Di :: 146Nd_TCRgd+PE
20.0%
Cd110Di :: 110Cd_CD4
Cd111Di :: 111Cd_CD8a
B
Flow cytometry gating
78.9%
FSC-H
FSC-A
51.8%
SSC-A
FSC-A
64.9%
CD45 AF700
Live/Dead 7AAD
67.3%
CD14 CD19 PacBlue
CD3 BUV805
43.4%
CD4 BV605
CD8 FITC

**Figure S4: Cytometry gating of CD8$^+$ T cells**
S4A: Representative gating of CD8$^+$ T cells in mass cytometry data
S4B: Representative gating of CD8$^+$ T cells in flow cytometry sorting data

**Table S1: Cohort demographics and clinical information**

**Table S2: Mass cytometry panel**

**Table S3: Mass cytometry peptide list**

**Table S4: Flow cytometry panel**

**Table S5: Antibody-derived tag (ADT) panel**
